# High-throughput determination of the antigen specificities of T cell receptors in single cells

**DOI:** 10.1101/457069

**Authors:** Shu-Qi Zhang, Ke-Yue Ma, Alexandra A. Schonnesen, Mingliang Zhang, Chenfeng He, Eric Sun, Chad M. Williams, Weiping Jia, Ning Jiang

## Abstract

We present tetramer-associated T-cell receptor sequencing (TetTCR-Seq), a method to link T cell receptor (TCR) sequences to their cognate antigens in single cells at high throughput. Binding is determined using a library of DNA-barcoded antigen tetramers that is rapidly generated by *in vitro* transcription and translation. We applied TetTCR-Seq to identify patterns in TCR cross-reactivity with cancer neo-antigens and to rapidly isolate neo-antigen-specific TCRs with no cross-reactivity to the wild-type antigen.

---

The ability to link T cell antigens, peptides bound by the major histocompatibility complex (pMHC), to T cell receptor (TCR) sequences is essential for monitoring and treating immune-related diseases. Fluorescently labeled T cell antigen oligomers, such as pMHC tetramers, are widely used to identify antigen-binding T cells^1^. However, spectral overlap limits the number of pMHC tetramer species that can be studies in parallel and the extent of cross-reactivity that can be examined^1^. CyTOF with isotope-labeled pMHC tetramers can interrogate a larger number of antigen species simultaneously, but its destructive nature prevents the linkage of antigens bound to TCR sequences^1^.

DNA-barcoded pMHC dextramer technology has been used for the analysis of antigen-binding T cell frequencies to samples of more than 1,000 pMHCs for T cells sorted in bulk^2^. However, with bulk analysis, information on the binding of single or multiple peptides to individual T cells is lost. In addition, antigen-linked TCR sequences cannot be obtained, which is valuable for tracking antigen-binding T cell lineages in disease settings, TCR-based therapeutics development^3^, and for uncovering patterns in TCR-antigen recognition^4^. Another limitation is the high-cost and long-duration associated with synthesizing peptides chemically^5^, which prevents the quick generation of a pMHC library that can be tailored to specific pathogens or neo-antigens in an individual.

To address these challenges, we developed TetTCR-Seq, for the high-throughput pairing of TCR sequences with potentially multiple pMHC species bound on single T cells. First, we constructed a large library of fluorescently labeled, DNA-barcoded (DNA-BC) pMHC tetramers in a rapid and inexpensive manner using *in vitro* transcription/translation (IVTT) (Fig. 1a). Next, tetramer-stained cells were single-cell sorted and the DNA-BC and TCRαβ genes were amplified by RT-PCR (Fig. 1b). A molecular identifier (MID) was included in the DNA-BC to provide absolute counting of the copy number for each species of tetramers bound to the cell. Finally, nucleotide-based cell barcodes were used to link multiple peptide specificities with their bound TCRαβ sequences (Fig. 1b). DNA-BC pMHC tetramers are compatible with isolation of rare antigen-binding precursor T cells^6^, making TetTCR-Seq a versatile platform to analyze both clonally expanded and precursor T cells.

**Figure 1:**
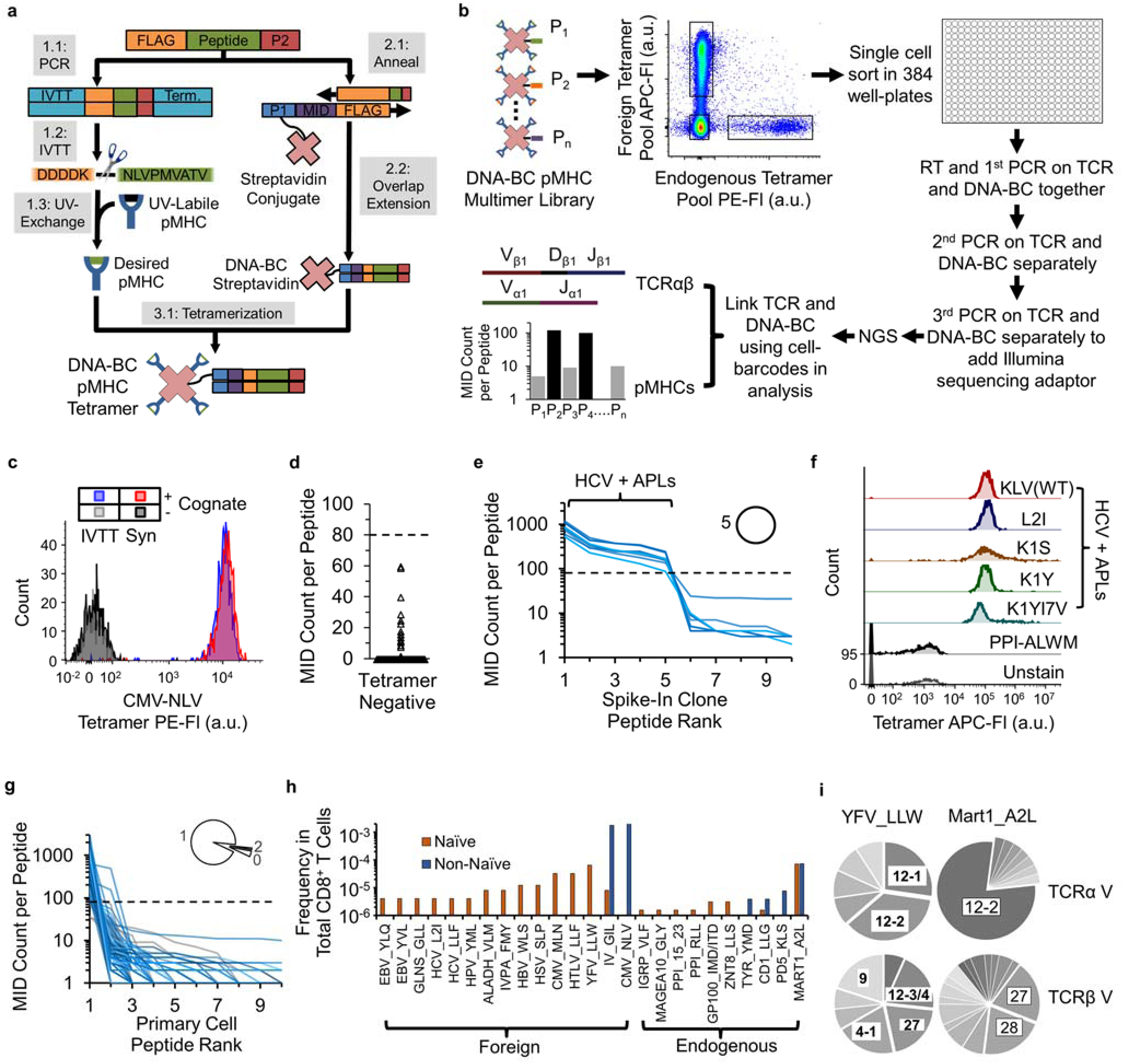
Workflow for generation of DNA-BC pMHC tetramer library and proof-of-concept of using TetTCR-Seq for high-throughput linking of antigen binding to TCR sequences for single T cells. **(a)** Workflow for generation of DNA-BC pMHC tetramers. **(b)** Workflow of TetTCR-Seq. **(c)** Comparison of staining performance for IVTT and synthetic peptide generated pMHC tetramers on T cell clones. Experiment repeated independently once with similar results. **(d)** MID counts per peptide detected on single T cells sorted from the Tetramer^-^ fraction in Experiment 1 (768 peptides from 8 cells). Dashed line, MID threshold. **(e)** Peptide rank curve by MID counts for each of top 10 ranked peptides for single sorted cells from the spike-in clone (8 cells) in Experiment 1. Dashed line is as in (d). Each blue solid line represents the MID counts associated with each of the 96 peptides that can potentially bind on a single cell. Inset, proportion of cells with the indicated number of positively binding peptides. **(f)** Fluorescent intensity of the HCV-KLV(WT) binding T cell clone, used as spike-in in Experiment 1, stained individually with the indicated pMHC tetramers in a separate experiment. Experiment performed once. **(g)** Peptide rank curve by MID counts as in (e) for the Tetramer^+^ primary T cell populations (167 cells) in Experiment 1. Grey solid lines indicate cells with no detected peptides. **(h)** Calculated frequencies of antigen-binding T cell populations in total CD8^+^ T cells for peptide with at least 1 detected T cell, separated by phenotype, in Experiment 1. **(i)** V-gene usage of unique TCR sequences that are specific for YFV_LLW (naïve and non-naïve combined, n = 11 cells for TRAV, n = 15 cells for TRBV) or MART1_A2L (naïve and non-naïve combined, n = 33 cells for TRAV, n = 43 cells for TRBV). P1, P2, and Pn, unique peptide ligands. NGS, next-generation sequencing. Fl, fluorescence intensity. a.u., arbitrary unit. APL, altered peptide ligand.

As constructing large pMHC libraries via UV-mediated peptide exchange using traditionally synthesized peptide is costly with long turnaround times^5^, we use a set of peptide-encoding oligonucleotides that serve as both the DNA-BCs for identifying antigen specificities and as DNA templates for peptide generation via IVTT (Fig. 1a). The IVTT step only adds a few hours once oligonucleotides are synthesized. This significantly reduces the cost (about 20-fold) and time (2-3 days instead of weeks) compared to synthesizing peptides chemically.

pMHC tetramers generated by our IVTT method have similar performance as the synthetic-peptide counterparts (Fig. 1c and Supplementary Fig. 1 and 2). DNA-BC conjugation did not interfere with staining and has comparable sensitivity to fluorescent readouts (Supplementary Fig. 3 and 4). Six main TetTCR-Seq experiments were performed (Supplementary Fig. 5). We first assessed the ability of TetTCR-Seq to accurately link TCRαβ sequences with pMHC binding from primary CD8^+^ T cells in human peripheral blood. In Experiment 1, we constructed a 96-peptide library consisting of well documented foreign and endogenous peptides bound to HLA-A2 (Supplementary Table) and isolated dominant pathogen-specific T cells as well as rare precursor antigen-binding T cells from a healthy CMV sero-positive donor (Fig. 1, Supplementary Fig. 6, 7). To test whether TetTCR-Seq can detect cross-reactive peptides, we included a documented HCV wildtype (WT) peptide, HCV-KLV(WT)^7^, and 4 candidate altered peptide ligands (APL) with 1-2 amino acid (AA) substitutions (Supplementary Table). An established HCV-KLV (WT) T cell clone^7^ was spiked into the donor’s sample to test its cross-reactive potential.

Bound peptides were classified by their MID counts using two criteria: an MID threshold derived from tetramer negative controls and a ratio of MID counts between the peptides above and below this threshold (Fig. 1d and Supplementary Information). Using this classification scheme, we identified the HCV-KLV(WT) epitope from all spike-in cells sorted (Fig. 1e, Supplementary Fig. 8a). In addition, we discovered that all four APLs were also classified as binders. A separate experiment confirmed the binding of these APLs to the T cell clone (Fig. 1f). These results show that TetTCR-Seq is able to resolve positively-bound peptides in primary T cells and identify up to five cross-reactive peptides per cell.

The majority of primary T cells were classified as binding one peptide (Fig. 1g). This result is expected because the probability of TCR cross-reactivity between similar peptides is higher than disparate ones^8, 9^. Among the peptides surveyed, we found a high degree of peptide diversity in the foreign-antigen-binding naïve T cells (Fig. 1h). This diversity reduced to two dominant peptides for CMV and influenza in the non-naïve repertoire^10^ (Fig. 1h). This is expected given the CMV sero-positive status and the high probability of influenza exposure or vaccination for this donor. The majority of cells within the endogenous-antigen-binding population bind MART1-A2L, which corroborates its high documented frequency^6, 10^ (Fig. 1h). Linked TCR and DNA-BC analysis revealed that TCRα V genes 12-2 and 12-1/12-2 dominate in MART1-A2L and YFV-LLW specific TCRs, respectively (Fig. 1i), which is consistent with other reports^11, 12^. TetTCR-Seq on a second CMV sero-positive donor (Experiment 2) verified the findings from Experiment 1 (Supplementary Fig. 9). These results highlight the ability of TetTCR-Seq to accurately link pMHC binding with TCR sequences.

Naïve T cells from healthy donors are a useful source of neo-antigen-binding TCRs^3^. However, most neo-antigens are 1 AA from the WT sequence, meaning that neo-antigen-binding TCRs can potentially cross-react with host cells to cause autoimmunity. Although clinical adverse effects caused by neo-antigen recognizing T cells cross-reacting with endogenous tissue have not been reported, possibly due to the lack of technology development, other forms of cross-reactivity have been reported to cause death in clinical trials^13^. We next applied TetATCR-Seq to study the extent of cancer antigen cross-reactivity in healthy donor peripheral blood T cells and isolate neo-antigen (Neo)-specific TCRs with no cross-reactivity to wildtype counterpart antigen (WT). In Experiment 3, we surveyed 20 pairs of Neo-WT peptides that bind with high affinity to HLA-A2. pMHC tetramer-based selection of naïve T cells has an inherent risk of selecting T cells reactive to peptides that are not naturally processed. As such, peptides were also chosen based on previous evidence of tumor expression and T cell targeting^3, 14-16^ (Supplementary Table). Neo and WT pMHC pools were labeled using two separate fluorophores, allowing sorting of three cell populations, Neo^+^WT^-^, Neo^-^WT^+^, and Neo^+^WT^+^ (Fig. 2a and Supplementary Fig. 10).

**Figure 2:**
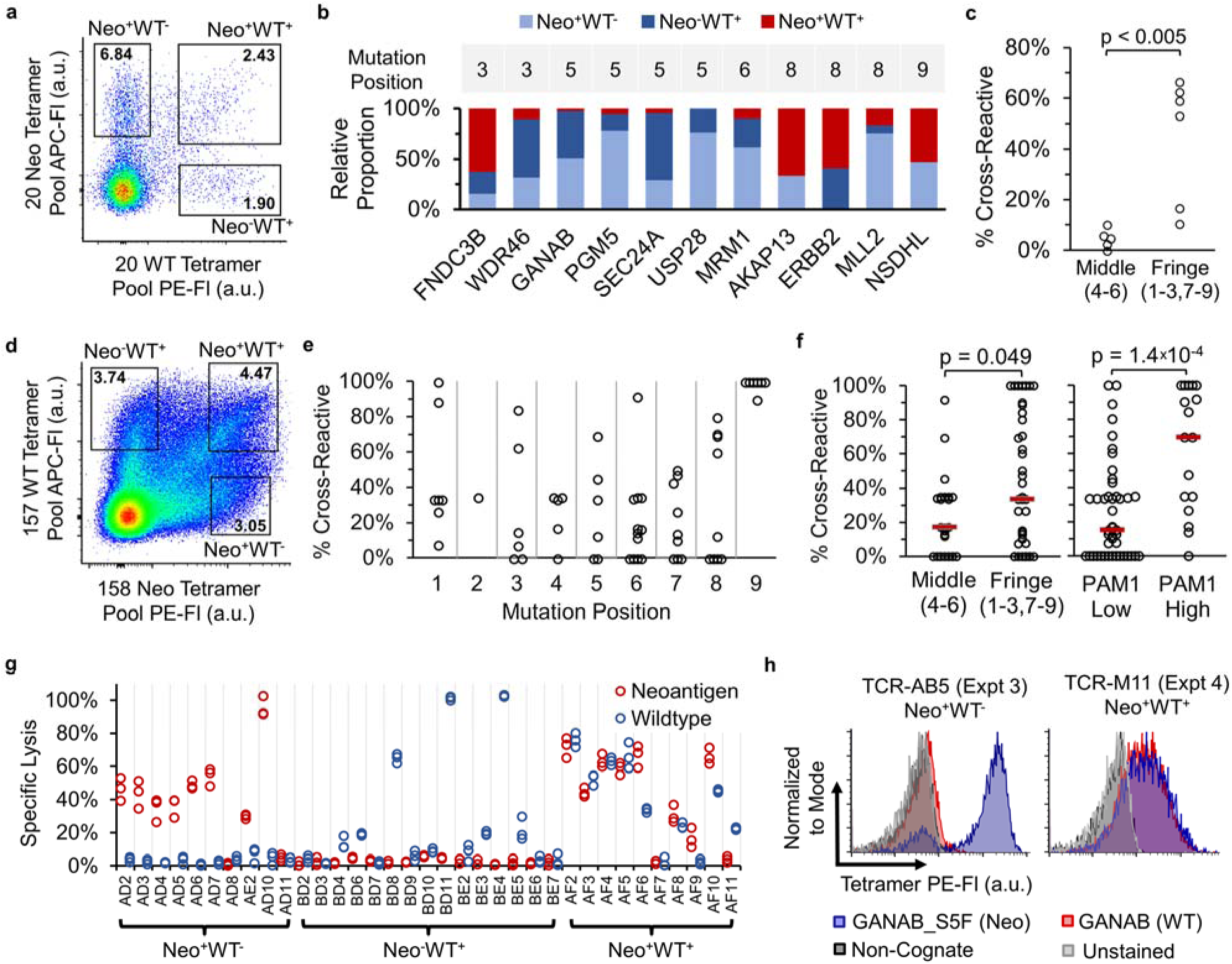
High prevalence of neo-antigen binding T cells that cross-react to WT counterpart peptides and high-throughput isolation of neo-antigen-specific TCRs for multiple specificities in parallel using TetTCR-Seq. (a-c) Experiment 3, isolation of single Neo and/or WT binding T cells from a healthy donor using a 40 Neo-WT antigen library. (a) DNA-BC pMHC tetramer staining profile of naïve CD8^+^ T cells from the tetramer pool-enriched fraction. See Supplementary Fig. 10 for gating scheme. (b) Relative proportion of T cells among the three possible antigen binding combinations (Neo^+^WT^-^, Neo^-^WT^+^, Neo^+^WT^+^) for each Neo-WT antigen-pair from Experiment 3. Only antigen-pairs where both peptides were detected in at least one cell and have at least three detected cells in total (149 cells, see Methods) were included. (c) Effect of neoantigen mutation position (indicated in parenthesis) on the proportion of cross reactive T cells from red bars in (b). (5 Neo-WT pairs for middle and 6 for fringe, One-tailed Mann Whitney U-Test). (d-f) Experiment 5 and 6, isolation of Neo and/or WT binding T cells using a 315 Neo-WT antigen library. (d) Staining profile as in (a) for Experiment 5. See Supplementary Fig. 15 for gating scheme. (e) Proportion of cross-reactive T cells for Neo-WT antigen-pairs based on mutation position. Same data filter as (b) is used. (62 Neo-WT pairs from 517 cells). (f) Effect of neoantigen mutation position as in (c) or PAM1 value on the proportion of cross reactive T cells in (e). Red bars denote median. (left-to-right, n = 23, 39, 45, 17 Neo-WT pairs, One-Tailed Mann Whitney U-Test). Alternative analysis using contingency tables are shown in Supplementary Figure 16. (g) LDH cytotoxicity assay on *in vitro* expanded primary T cell lines sorted using DNA-BC pMHC tetramers as in (a) interacting with T2 cells pulsed with the 20 neo-antigen peptide pool or 20 WT counterpart peptide pool. Each condition was performed in triplicates derived from separate wells of cells. (h) Staining of Jurkat 76 cell line transduced with TCRs from Experiment 3 and 4 with the indicated tetramers. Experiment was performed once. Based on TetTCR-Seq of the original T cells, TCR-AB5 recognized the neo-antigen GANAB_S5F while TCR-M11 recognized both GANAB_S5F and its WT counterpart, GANAB.

T cells with two detected peptide binders accounted for 84% of the Neo^+^WT^+^ population, 98% of which belonged to a Neo-WT antigen-pair (Supplementary Fig. 11). Cells in the Neo^+^WT^+^ population bound 11 of the 20 Neo-WT antigen-pairs, indicating that Neo-WT cross-reactivity is wide-spread in the precursor T cell repertoire (Fig. 2b). By analyzing the proportion of mono- and cross-reactive T cells from each Neo-WT pair, we observed that neo-antigens with mutations at fringe positions 3, 8, and 9 elicited significantly more cross-reactive T cells than the ones at center positions 4, 5, and 6 (Fig. 2c). This is consistent with observations made by others using alanine substitutions on peptides in a mouse model^17^. TetTCR-Seq on a separate donor (Experiment 4) showed the same trend (Supplementary Fig. 13), indicating that this property is conserved between donors for the peptides tested.

To test the feasibility of TetTCR-Seq to screen larger libraries, we assembled a 315 Neo-WT antigen-pair library and profiled T cell cross-reactivity in more than 1,000 Tetramer^+^ CD8^+^ sorted single T cells from two donors (Experiment 5 and 6, Fig. 2d, and Supplementary Fig. 15,16). ELISA on all 315 pMHC species showed no difference in pMHC UV-exchange efficiency between detected and undetected peptides (Supplementary Fig. 17). Similar to Experiment 3 and 4, neo-antigen mutations in the fringes have elevated percentages of cross-reactive T cells than mutations in the middle (Fig. 2e,f, Supplementary Fig. 16j). Using this larger dataset, we also found that neo-antigen mutations with high PAM1 values, a surrogate for chemical similarity related to evolutionary mutational probability^18^, have a significantly higher percentage of cross-reactive T cells than those with low PAM1 values (Fig. 2f, ). Thus, in addition to mutation position, WT-binding T cells are more likely to recognize the neo-antigen if the mutated AA is chemically similar to the original. Additional, so far unaccounted variations still exist between peptides, highlighting the necessity for experimental screening against WT cross-reactivity when using neo-antigen based therapy in cancer.

We also assessed the utility of TetTCR-Seq to isolate neo-antigen-specific TCRs with no cross-reactivity to WT. We generated primary cell lines, each derived from 5 sorted cells in the Neo^+^WT^-^, Neo^-^WT^+^, and Neo^+^WT^+^ populations from Experiment 3 and 4. These cells lysed antigen-pulsed target cells in a manner that matched their gating scheme during sorting, independent of the choice of pMHC tetramer fluorophore (Fig. 2g). Further TetTCR-Seq analysis of Neo^+^WT^-^ and Neo^+^WT^+^ T cell lines showed unique TCRs in each cell line targeting a limited range of antigens (Supplementary Fig. 19a and b). Cytotoxicity test confirmed the cross-reactivity of Neo^+^WT^+^ cell lines as identified by TetTCR-Seq (Supplementary Fig. 19c). Lastly, the antigen-recognition of Jurkat76 cell lines transduced with TCRs identified from Experiment 3 and 4 matched their original specificities (Fig. 2h, Supplementary Fig. 20). Together, our T cell line and TCR-transduced Jurkat experiments show that TetTCR-Seq is not only capable of identifying cross-reactive TCRs on a large scale but can also identify functionally reactive neo-antigen specific TCRs that are not cross-reactive to WT-peptide in a high-throughput manner, which could be valuable in TCR re-directed adoptive cell transfer therapy^3, 19^.

In conclusion, we show that TetTCR-Seq can accurately link TCR sequences with multiple antigenic pMHC binders in a high-throughput manner, which can be broadly applied to interrogate antigen-binding T cells in T cell populations, from infection to autoimmune disease and cancer immunotherapy, potentially even for individual patients. With methods emerging for predicting antigenic pMHCs for groups of TCR sequences^4^, TetTCR-Seq can not only facilitate development in this area but also help to validate informatically predicted antigens. Lastly, TetTCR-Seq can be integrated with single-cell transcriptomics and proteomics to gain further insights into the connections between single T cell phenotype, and TCR sequence and pMHC-binding landscape ^20^.

## Acknowledgements

We thank Dr. Ben Wendel for helpful discussions and for producing recombinant HLA-A2; Dr. Mark M. Davis and Dr. Huang Huang at Stanford University for helpful discussion of the lentiviral transduction protocol and providing a template TCR construct and HLA-A2 construct; Dr. Wolfgang Uckert at Max Delbruck Center for Molecular Medicine for sharing the Jurkat 76 cell line; Dr. Amy Brock at UT Austin for sharing HEK 293T cell line; Dr. Jizhong Lou at the Institute of Biophysics, Chinese Academy of Sciences, for helping with HCV APL prediction; Patrik Parker, Kushal Patel, and Henry Pan for assistance with initial prototyping; the NIH tetramer center for additional pMHC tetramer reagents. We also thank anonymous blood donors and staff members at We Are Blood for sample collection. This work was supported by NIH Grants R00AG040149 (N.J.), S10OD020072 (N.J.), R33CA225539 (N.J), by NSF CAREER Award 1653866 (N.J.), by the Welch Foundation Grant F1785 (N.J.), by Robert J. Kleberg, Jr. and Helen C. Kleberg Foundation (N.J.), and by National Natural Science Foundation of China major international (regional) joint research project 81220108006 (W.J.) and NSFC-NHMRC joint research grant 81561128016 (W.J). N.J. is a Cancer Prevention and Research Institute of Texas (CPRIT) Scholar and a Damon Runyon-Rachleff Innovator. S.-Q.Z. is a recipient of Thrust 2000—Archie W. Straiton Endowed Graduate Fellowship in Engineering No. 1. A.A.S. is a recipient of the Cockrell School of Engineering fellowship and the Thrust 2000 - Mario E. Ramirez Endowed Graduate Fellowship in Engineering.

## Author Contributions

S.-Q.Z. conceived and developed the technology platform. S.-Q.Z and N.J. conceived and designed the study. S.-Q.Z. and K.-Y.M. designed, performed, and analyzed data for the majority of experiments; A.A.S. and M.Z. performed TCR cloning, transduction, and pMHC tetramer staining studies; C.H. wrote the script for converting sequencing data into TCR sequences, DNA-BC, and MIDs, and predicted HCV APLs; C.M.W. and E.S. performed in vitro cell culture and functionality experiments; W.J. co-supervised study and co-designed some experiments; N.J. supervised the study; S.-Q.Z. and N.J. wrote the manuscript with feedback from all authors.

